# Construction and analysis of an interologous protein-protein interaction network of *Camellia sinensis* leaf (TeaLIPIN) from RNA-Seq datasets

**DOI:** 10.1101/592394

**Authors:** Gagandeep Singh, Vikram Singh, Vikram Singh

**Affiliations:** Centre for Computational Biology and Bioinformatics, School of Life Sciences, Central University of Himachal Pradesh, Dharamshala, India-176206

**Keywords:** *Camellia sinensis*(Tea), PPI network, RNA-Seq data, Leaf transcriptome, Interolog, KEGG pathways, Transcription factors (TFs)

## Abstract

Worldwide, tea (*Camellia sinensis*) is the most consumed beverage primarily due to the taste, flavour and aroma of its newly formed leaves; and has been used as an important ingredient in several traditional medicinal systems because of its antioxidant properties. For this medicinally and commercially important plant, design principles of gene-regulatory and protein-protein interaction networks at sub-cellular level are largely un-characterized mainly because neither its reference assembly nor annotated genome is available though plenty of its transcriptomes have been studied. In this work, we report a tea leaf interologous PPI network (TeaLIPIN) consisting of 11,208 nodes and 1,97,820 interactions. A reference transcriptome assembly was first developed from all the 44 samples of 6 publicly available leaf transcriptomes (1,567,288,290 raw reads). By inferring the high-confidence interactions among potential proteins coded by these transcripts using known experimental information about PPIs in 14 plants, an interologous PPI network was constructed and its modular architecture was explored. Comparing this PPI network with 10,000 realisations of corresponding random networks (Erdős-Rényi models) and examining over three network centrality metrics, we predict 2,931 bottleneck proteins (having *p-values* <0.01). 270 of these are deduced to have transcription factor domains by developing the HMM models of known plant TFs. Final transcripts were also mapped to the draft tea genome in order to search the probable loci of their origin. We believe that the proposed novel methodology can easily be adopted to develop and explore the PPI interactomes in other plant species by making use of the available transcriptomic data.

## Introduction

*Camellia sinensis* is an important commercial crop that is consumed worldwide as the most popular non-alcoholic beverage and has also been used as a traditional medicine in various parts of the world since c. 3000 BC (Xia et al. 2017). Its leaves are known to have several anti-oxidants, anti-cancerous and anti-inflammatory properties as it contains wide range of bioactive ingredients having important roles in treatment of several diseases (Namita et al. 2012). Due to several microbial attacks and environmental stresses, various processes related to the growth and developments of tea leaves are continuously affected that severely hamper the quality and production of this crop (Zhou et al. 2017). To understand the molecular insights of these processes in detail and to explore the novel strategies for the potential control of these processes, it is crucial to identify key proteins and their interactions with others that directly or indirectly regulate the expression of various genes that in-turn regulate primary and secondary metabolic pathways etc. (Xia et al. 2017). Although, for understanding the architecture of genetic circuitry of tea, a deluge of information has been generated by means of genome or transcriptome sequencing and several thousands of genes, their potential proteins with functions have been predicted; to the best of our knowledge till date no attempt has been made towards studying regulatory mechanisms of action at molecular level by developing and examining the global interactomes.

Regulation of various biological processes is governed by a web of protein-protein interactions at molecular level (Jeong et al. 2001). PPI networks form the basis of various regulatory and signalling processes such as signal transduction, stress response, homeostasis, plant defense, and organ formation at cellular level etc. (Zhang et al, 2007; Geisler-Lee et al, 2007). Interologous PPI networks have been widely studied for the model crops such as *Arabidopsis thaliana* (Geisler-Lee et al, 2007), *Oryza sativa* (Zhu et al. 2011), *Zea mayz* (Zhu et al. 2016) and *Manihot esculenta* (Thanasomboon et al. 2017) to address various biological questions. Interolog based method to construct a PPI network has been reported to be useful in the cases where reference assembly or annotated genome was unavailable (Yu et al, 2004). In cases, where reference assembly or annotated genome is not available for a species, it is difficult to identify or predict PPI interactions and to study the interactions at the systems level. To address that research gap, we propose a methodology to develop an interolog based PPI network by using the publicly available transcriptomes in order to highlight the regulatory behaviours of various proteins among each other.

In this work, we have selected the leaf transcriptomic data of *Camellia sinensis* available in the public domain from 6 different studies related to biotic and abiotic stresses to identify the potential candidates involved in various processes related to several regulatory pathways. A reference transcriptome assembly was first developed from all the experiments followed by identification of interactions among translated proteins across various plant species to construct an interologous PPI network. We also have mapped the predicted nodes on the draft tea genome. This purposed PPI network provides a bunch of several possible interactions that will be helpful in addressing various research queries towards finding novel proteins that regulate important mechanisms related to growth and development of tea. The overall flow-chart of our work is shown in Fig. 1.

**Fig. 1.**
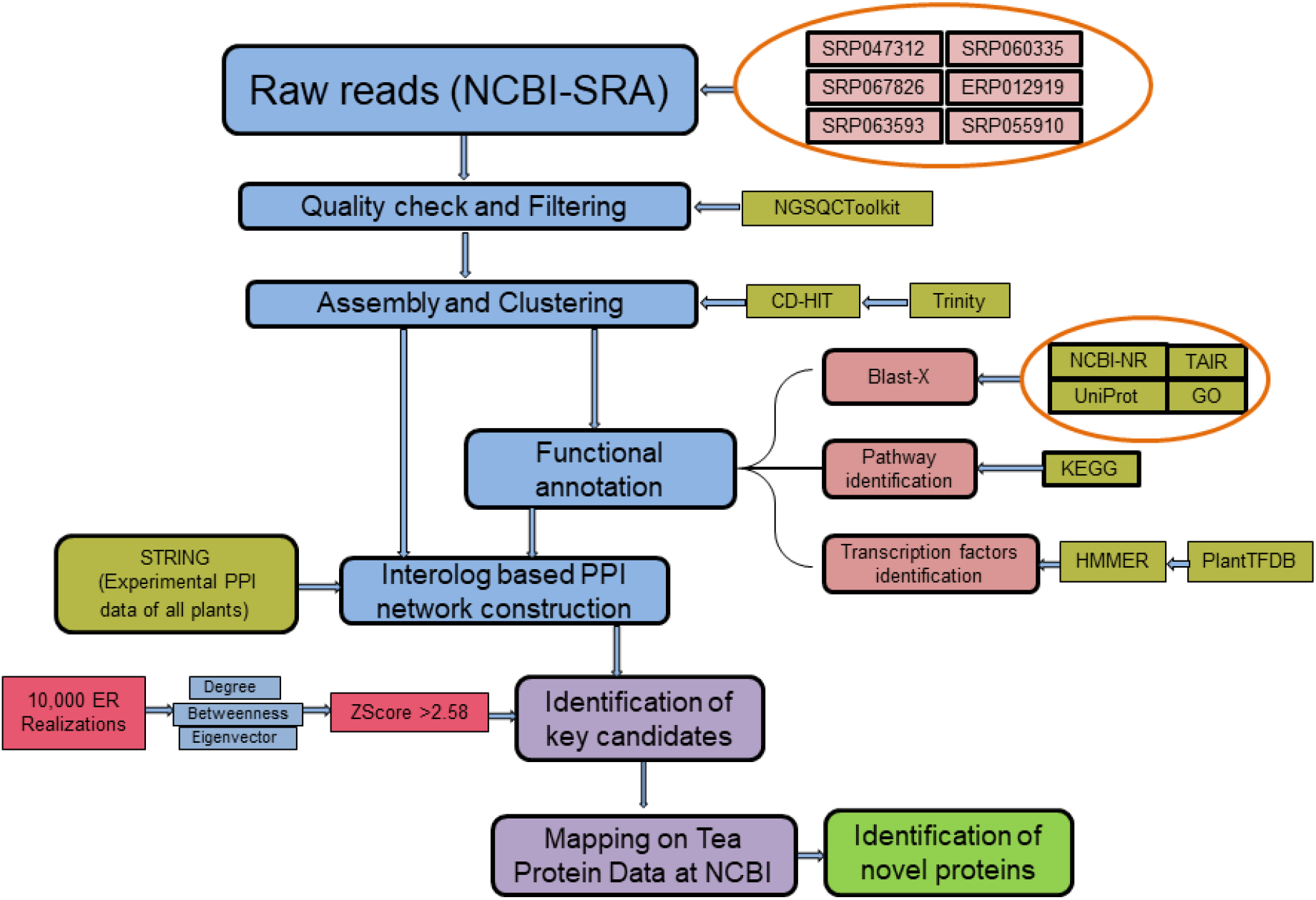
Work flow chart representing various steps involved in construction and analysis of TeaLIPIN.

## Materials and Methods

### Data mining and assembly

Paired-end raw reads from 6 leaf transcriptomes of *Camellia sinensis* were downloaded from NCBI sequence read archive database (https://www.ncbi.nlm.nih.gov/) with average read lengths of 33, 72, 100, 100, 101 and 151 from six experiments having SRA accessions SRP047312 (Paul et al. 2014), SRP067826 (Jayaswall et al. 2016), SRP060335 (Shi et al. 2013), ERP012919 (Zhang et al. 2017), SRP063593 (Li et al. 2018) and SRP055910 (Li et al. 2016), respectively. All the raw reads were filtered separately (experiment-wise) based on the average read length using NGS QC toolkit by setting 70% cut-off read length and cut-off quality score of 20 for obtaining the high quality reads (Patel and Jain 2012). Next, all the six datasets were assembled separately using Trinity (version r20140717) with default parameters (Grabherr et al. 2011). Finally, to remove redundancy, all the six assembled datasets were subjected to clustering with CD-Hit with a cut-off of 90% similarity (Li and Godzik 2006). Final dataset consisting of 2,61,695 transcripts was used in the further analysis.

### Functional annotation of assembled transcripts

Functional annotations to the final assembled transcripts were assigned by subjecting them to the BLASTX against non-redundant (NR) database at NCBI (Altschul et al. 1990), UniProt (Apweiler et al. 2004) and TAIR database (Berardini et al. 2015). Gene Ontology annotation for the functionally identified genes was performed by AgriGo (Du et al. 2010) and WEGO (Ye et al. 2006) tools. Pathways mapping was performed through KEGG database (Kanehisa et al. 2016). We attempted to computationally characterize the tea leaf transcripts that may have the transcription factor like activities, as a case study in this work. From each of the 58 categories of transcription factors (TFs) in plant transcription factor database (Jin et al. 2016), sequences of all the TFs were retrieved and their alignment was obtained using standalone version of Clustal Omega (Siever et al. 2011). Hidden Markov models of TFs from each category were built using HMMER (http://hmmer.janelia.org/). Largest open reading frames (ORFs) were searched in the final assembled transcripts and these ORFs were then subjected to the HMMs for predicting if a transcript may belong to any of the 58 categories of the plant transcription factors.

### Interolog based tea leaf protein-protein interaction network (TeaLIPIN) construction

Experimentally validated pre-determined protein-protein interactions in all the 14 plant species (*Arabidopsis thaliana, Brachypodiumdistachyon, Brassica rapa, Glycine max, Hordeum vulgare, Musa acuminata, Oryza sativa, Populustrichocarpa, Sorghum bicolor, Setariaitalica, Solanum lycopersicum, Selaginellamoellendorffii, Vitis vinifera and Zea mays*) were obtained from STRING database, as available in May, 2018 (Szklarczyk et al. 2014) and the proteomes of all these 14 plants were downloaded from UniProt database (http://www.uniprot.org/). To find the high confidence orthologs of the assembled tea leaf transcripts, their BLASTX search was carried out against these plant proteomes using highly stringent parameters (sequence identity > 60%, sequence coverage > 80% and e-value ≤ 10^−10^) (Altschul et al. 1990). An interologous PPI network of tea leaf was created by mapping the obtained orthologs of tea transcripts to the experimental PPIs obtained via STRING. An interaction between a pair of tea transcripts was considered to be true, if there existed at least one interaction between their corresponding orthologs (Thanasomboon et al. 2017). Final constructed network (TeaLIPIN) was visualized and analyzed by Cytoscape v3.3.0 (Smoot et al. 2010). Final assembled transcripts appearing as a node in the TeaLIPIN that were potentially encoding the transcription factors were further mapped on the available draft genome of tea (Xia et al. 2017).

### Network modularity and metrics analysis

In order to search the presence of functional modules in the final network (TeaLIPIN), it was subjected to clustering using MCODE (Bader and Houge 2003). All the proteins present in the top 10 clusters (sorted on the basis of node degree) were analysed for pathway enrichment using DAVID Bioinformatics Resourses v6.8 (Huang et al. 2008). The largest component of TeaLIPIN was considered for the further analysis since several proteins were lying as outgroups, which may be due to an artefact of non-availability of full proteomic information in tea. Three network centralities (degree, betweenness and eigenvector) were computed for the TeaLIPIN and its random realizations. For a graph *G* (*V*, *E*) with *V* nodes and *E* links, *A* = (*a_i,j_*) represents the adjacency matrix of *G* and (*a_i,j_*) = 1 if there exists an edge between *i, j* and 0 otherwise. Degree centrality (*C_d_*) of a node *v* is given by the number of directly connected nodes and is defined as *C_d_*(*v*) = Σ_j_*A*(*v*,*j*). Greater the degree of a node, higher the importance it has since it can regulate more number of proteins and hence represent key a node. PPI networks follow power-law in their degree distribution and the higher degree nodes (hubs) tend to be more central in the network (Jeong et al. 2001).

Betweenness centrality (*C_b_*) of a node represents its importance to act as a connecting link between other nodes (Joy et al. 2005). For any two nodes *i, j* | *v* ∈ *V*, if the total number of paths existing between *i, j* is given by *σ_i, j_* and *σ_i, j_*(*v*) represents the number of paths going through node *v*, then *C_b_* of a node *v* is defined as 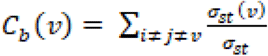. Eigenvector centrality (*C_ev_*) prioritizes the nodes in a network by considering the influence of its immediate neighbours also along with the number of connections it has (Newman 2008).

### ER network construction and significance evaluation to identify key proteins

*G_n,m_* type ER model based random networks that preserve size and average degree of a given network were constructed corresponding to the largest component of TeaLIPIN (Erdős and Rényi 1960). We constructed 10,000 such random realizations and computed above mentioned three centrality measures for the TeaLIPIN and each random network that were further used to calculate the significance score of each node. Statistically significant bottlenecks (high scoring nodes) were identified using z-score which is defined as 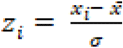, where 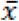 is the mean and *σ* is the standard deviation of random ensemble. All the nodes of TeaLIPIN with a p_value_ < 0.01 were termed as the key nodes. The proteins corresponding to these key nodes were further analyzed for their involvements in various pathways and their potential to code for a transcription factor.

## Results and Discussions

### Data mining and analysis

Total 1,567,288,290 (~1.5 Billion) paired-end reads were downloaded from publicly available NCBI-SRA database of six tea leaf transcriptomes. These transcriptomic experiments have been reported for the studies of three abiotic stresses, namely, cold stress (SRP047312), salinity and drought stress (ERP012919) and environmental stresses on leaves (SRP055910); and three biotic stresses, namely, fungal infection (SRP067826), mechanical wounding (SRP063593) and methyl jasmonate treatment (SRP060335). In each experimental data, quality checking and filtering of the raw reads was performed, and a total of 1,439,994,212 high quality cleaned reads were obtained. Since the read lengths were not uniform, these cleaned reads were first assembled separately for each experimental data. A total of 26,711; 64,034; 108,656; 64,458; 72,581 and 164,159 assembled sequences were obtained for SRP047312, SRP067826, SRP060335, ERP012919, SRP063593 and SRP055910 datasets respectively. Now to remove the redundancy in the entire data, all the assembled reads were subjected to stringent clustering that resulted in 2,61,695 final transcripts (S1 Supplementary File) with N50 value of 1,219 and average transcript length of 915. Details about the entire data are given in Table 1.

**Table 1.**
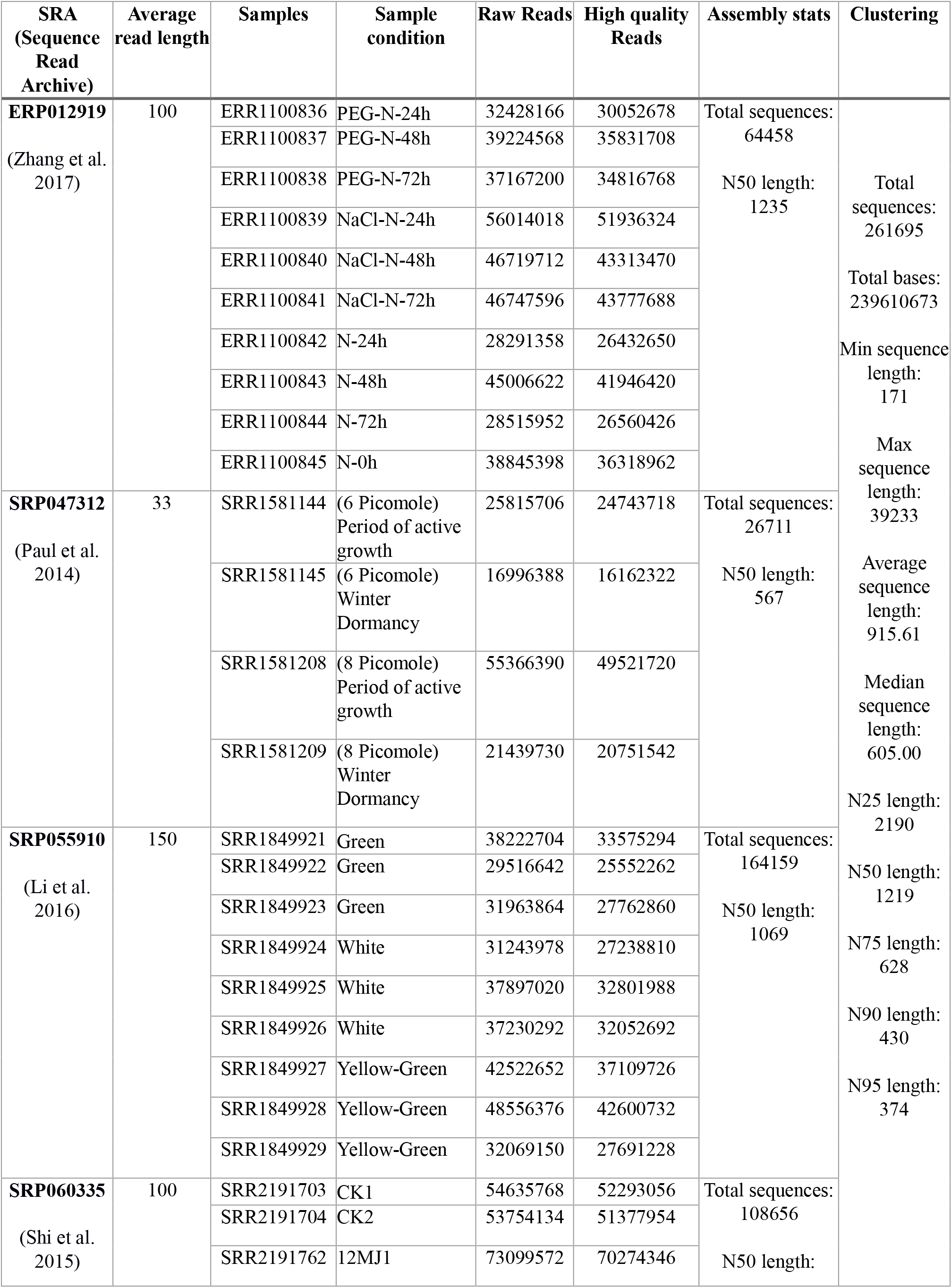
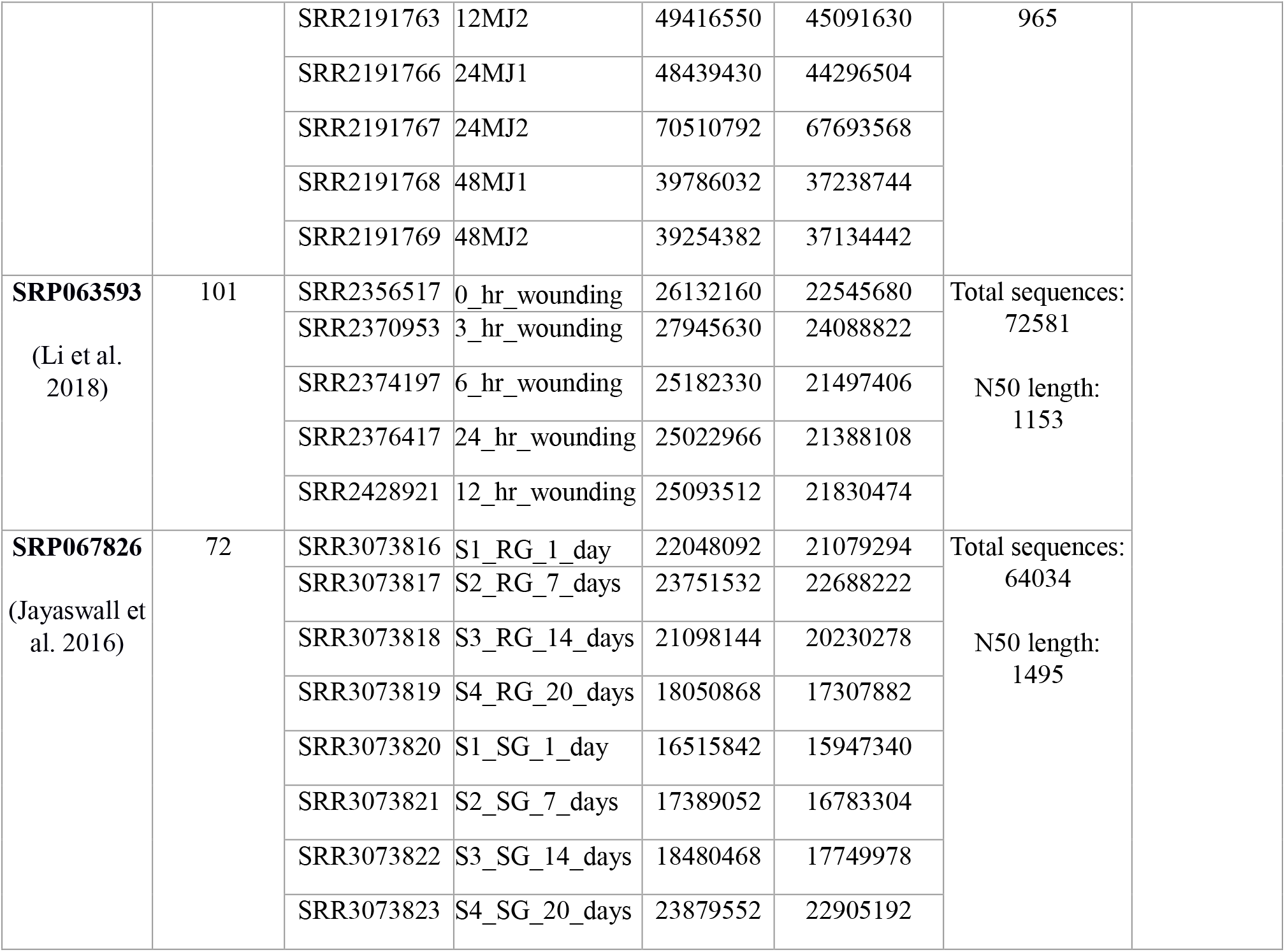
Details of the SRA samples selected for the current study.

### Functional annotation of assembled transcripts

In order to elucidate the function of final transcripts, three-fold comparative homology was performed against NRDB (at NCBI), TAIR and UniProt. We obtained 1,17,242 hits with non-redundant database at NCBI, 93,077 hits from TAIR and 84,123 hits from UniProt database, that resulted in a total of 1,19,178 mappings to the annotated proteins [S2 Supplementary File]. By gene ontology annotation, 51,947 GO terms were successfully predicted for 69,281 of mapped transcripts. Among these GO terms, about ~31% were biological processes classified into 37 categories, ~30% were cellular components divided into 21 functional categories and ~39% were molecular functions further classified into 36 categories. In biological processes, ‘cellular process and metabolic process’ were found to be most enriched followed by ‘response to stimulus’ and ‘biological regulation’. In cellular components, ‘cell’ and ‘cell part’ were highly enriched followed by ‘organelle and membrane’ and ‘organelle part protein-containing complex’. ‘catalytic activities and binding’ was the most enriched category in molecular functions followed by ‘transcriptional regulator activity’, ‘transporter activity’ and ‘structural molecular activity’ (Fig. 2). Assignment of gene ontology (GO) terms to several transcripts reveals the presence of diverse gene families in *Camellia sinensis*.

**Fig. 2.**
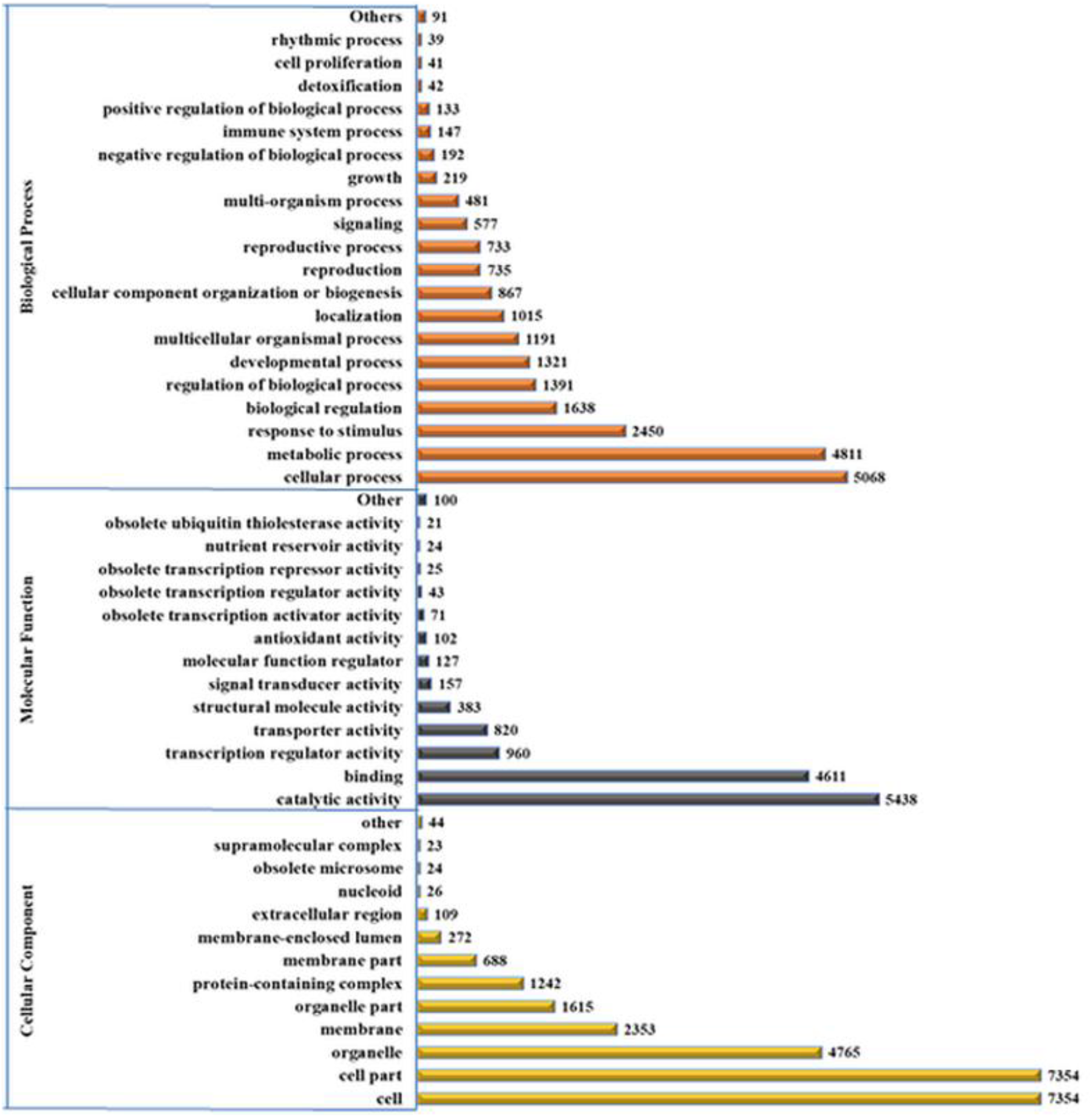
Gene ontology (GO) annotation of all the final assembled transcripts categorized in cellular component, biological process and molecular functions.

To identify the important regulatory pathways, in which our final transcripts might be involved in, all the 2,61,695 transcripts were mapped to KEGG database. Only 13,273 transcripts could be successfully mapped and predicted to have role in 416 pathways. These pathways were further classified into metabolism, genetic information processing, environmental information processing, cellular processes and organismal systems as shown in Fig. 3. In metabolism category, 11 metabolic pathways were predicted in which highest number of transcripts were found in carbohydrate metabolism (1,034 transcripts) followed by energy metabolism (780 transcripts) and amino acid metabolism (647 transcripts). In genetic information processing, 4 categories were found with 1,630 transcripts in translation category, 922 transcripts in folding, sorting and degradation category, followed by 417 transcripts in transcription category and 250 transcripts in and replication and repair category. Under environmental information processing, 3 categories were found with highest number of transcripts in signal transduction category (1,339 transcripts) followed by membrane transport and signaling molecules and interaction category with 54 and 30 transcripts respectively. In cellular processes, 4 categories were found that are transport and catabolism, cell growth and death, cellular community and cell motility involving 858, 630, 352 and 116 transcripts respectively. Finally, in the organismal systems, 10 categories were predicted with highest number of transcripts in environmental adaption (610 transcripts) followed by endocrine systems (549 transcripts) and immune system (459 transcripts).

**Fig. 3.**
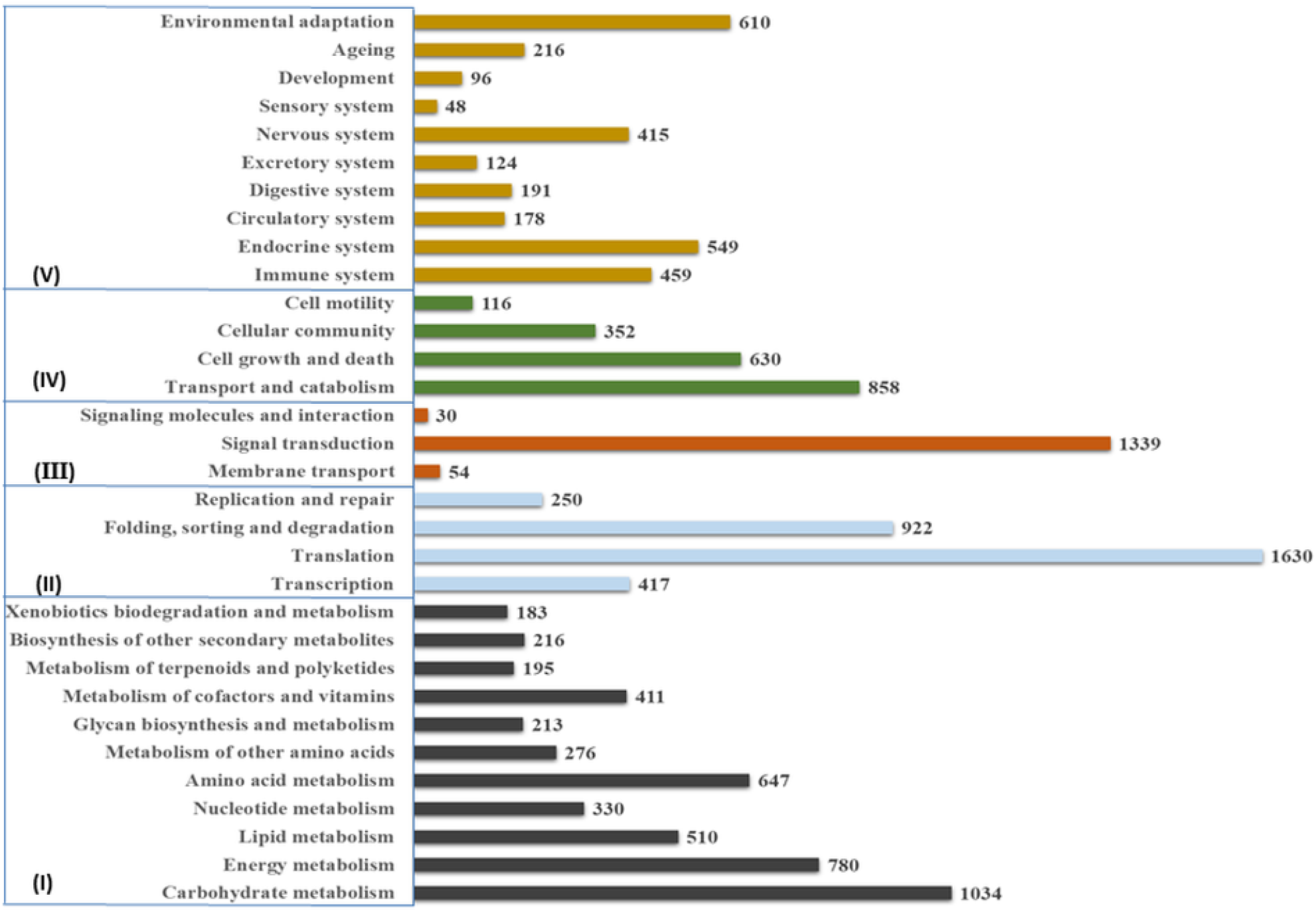
Pathwaysclassification of the final assembled transcripts into five major categories, I: Metabolism, II: Genetic information processing, III: Environmental information processing, IV: Cellular processes and V: Organismal systems.

Since transcription factors (TFs) are known to have very specific role in signaling and various other cellular processes during the stress responses, as a special case study we attempted to characterize the tea TFs. For that, all the 2,61,695 transcripts were screened to search for longest ORFs and 2,55,650 ORFs were found. The transcripts having ORFs were then subjected to the in-house developed 58 HMMs corresponding to these many transcription factor families of plants. 26,307 transcripts were resulted by the HMMs as the transcription factors and were spanning across all the 58 transcription factor families. The largest category of transcription factor is WRKY with 2,467 transcripts followed by 2,319 transcripts in S1Fa-like transcription family. Other, transcription factors like bHLH, STAT, MYB_related, E2F/DP, NAC have 1,790; 1,716; 1,355; 1,267; 1063 transcripts respectively as shown in Fig. 4.

**Fig. 4.**
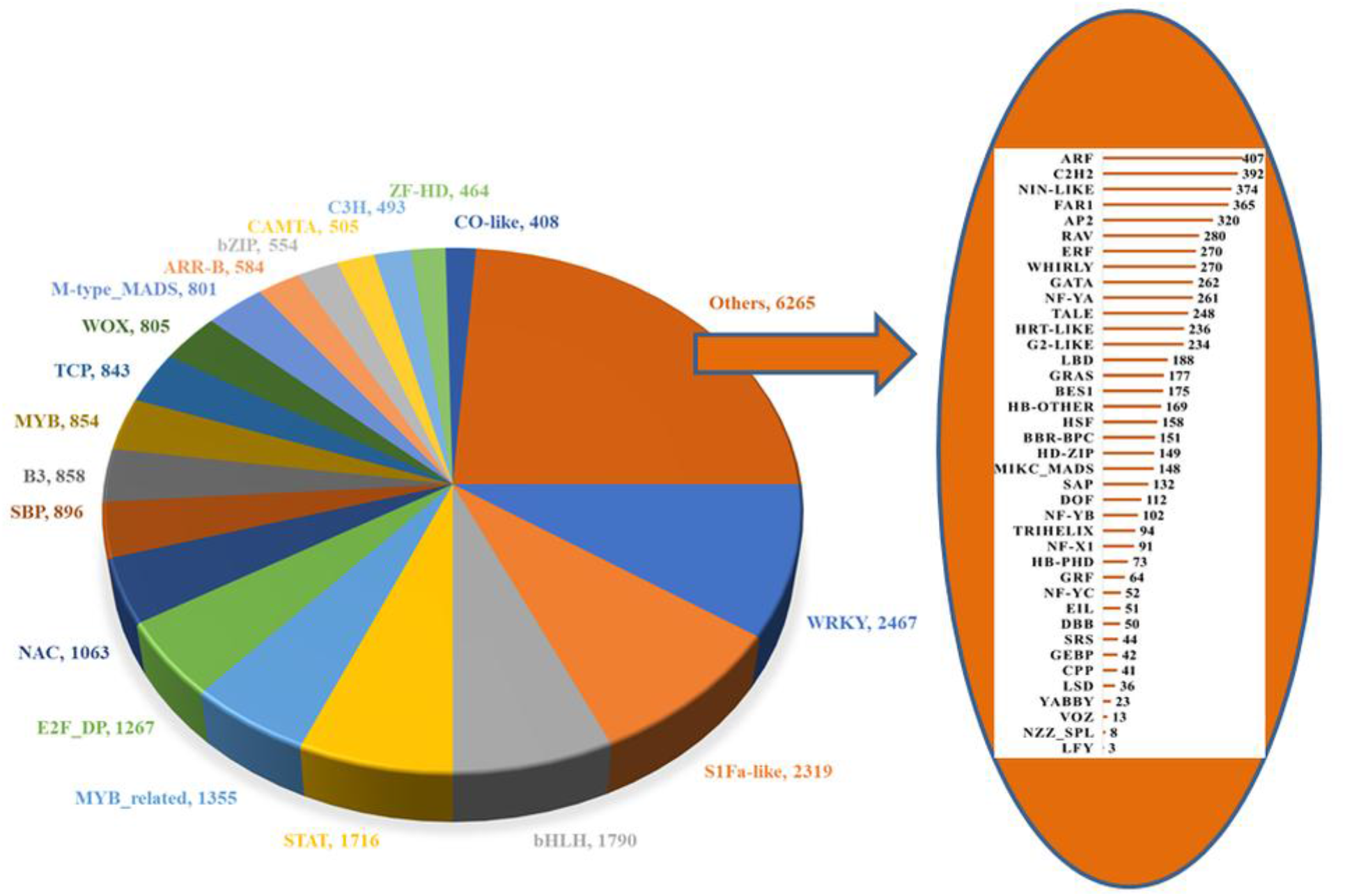
Characterization of transcription factors (TFs) identified from final assembled transcripts into 58 TF families.

### Construction of tea leaf interologous protein-protein interaction network (TeaLIPIN)

Since very little information about the protein-protein interactions in the tea proteome is known, experimentally validated interactions in pre-determined networks of 14 plant species from STRING database were selected as templates. Proteomic information about all these selected templates was first extracted from UniProt database and all the final assembled 2,61,695 transcripts were mapped to these templates via BLASTX with e-value 10^−10^. The 14 plants that were used in this step with the number of hits found are *Arabidopsis thaliana:* 87,762; *Brachypodium distachyon:* 81,097; *Brassica rapa:* 82,958, *Glycine max:* 88,639; *Hordeum vulgare:* 73,180; *Musa acuminate:* 78,095; *Oryza sativa:* 80,499; *Populus trichocarpa:* 88,344; *Sorghum bicolor:* 80,451; *Setaria italica:* 74,440; *Solanum lycopersicum:* 87,898; *Selaginella moellendorffii:* 61,488; *Vitis vinifera:* 95,784 and *Zea mays:* 81,779.

All the obtained orthologs from first species were selected and the novel unique interactions from subsequent species were added to this pool to generate the TeaLIPIN. Details about the relative contributions of various plant species in the construction of this network are given in Table 2. A total of 11,208 nodes with 1,97,820 interactions are successfully predicted [S3 Supplementary File] using this interolog based approach [Fig. 5 (a)]. We would like to remark here that a particular ordering of various species while generating the network will not have any impact on the final outcome, since these are the unique interactions in a species that are contributing in the build-up of the TeaLIPIN. Also, as this network has been constructed using a very large repository of experimentally validated interactions, it is highly unlikely that by adding or subtracting one or two species will have any substantial impact on the overall global structure of the TeaLIPIN.

**Table 2.** Details of the 14 plants selected as template for identifying the orthologous interaction pairs and their contribution in the construction of TeaLIPIN.

| Sr. No. | Plants | Experimental interactions | Interolog interactions obtained | Interactions added in the TeaLIPIN (%) | Cummulative interactions in TeaLIPIN |
| --- | --- | --- | --- | --- | --- |
| 1. | *Arabidopsis thaliana* | 72,709 | 1,583 | 1,583 (100%) | 1,583 |
| 2. | *Brachypodium distachyon* | 60,534 | 15,943 | 15892 (99.7%) | 17,475 |
| 3. | *Brassica rapa* | 204,493 | 43,461 | 41701 (95.5%) | 59,176 |
| 4. | *Glycine max* | 232,084 | 28,914 | 25093 (86.7%) | 84,269 |
| 5. | *Hordeum vulgare* | 40,686 | 1,420 | 1203 (84.7%) | 85,472 |
| 6. | *Musa acuminata* | 173,846 | 40,680 | 33907 (83.3%) | 119,379 |
| 7. | *Populus trichocarpa* | 150,119 | 25,441 | 18186 (71.4%) | 137,565 |
| 8. | *Oryza sativa* | 89,054 | 19,976 | 13118 (65.6%) | 150,683 |
| 9. | *Sorghum bicolor* | 72,312 | 9,557 | 4924 (51.52%) | 155,607 |
| 10. | *Setaria italica* | 65,138 | 20,251 | 9691 (47.8%) | 165,298 |
| 11. | *Zea mays* | 226,835 | 6,413 | 3049 (47.5%) | 168,347 |
| 12. | *Solanum lycopersicum* | 99,191 | 27,942 | 13207 (47.2%) | 181,554 |
| 13. | *Selaginella moellendorffii* | 98,857 | 15,663 | 7185 (45.8%) | 188,739 |
| 14. | *Vitis vinifera* | 69,832 | 21,322 | 9081 (42.5%) | 197,820 |

**Fig. 5.**
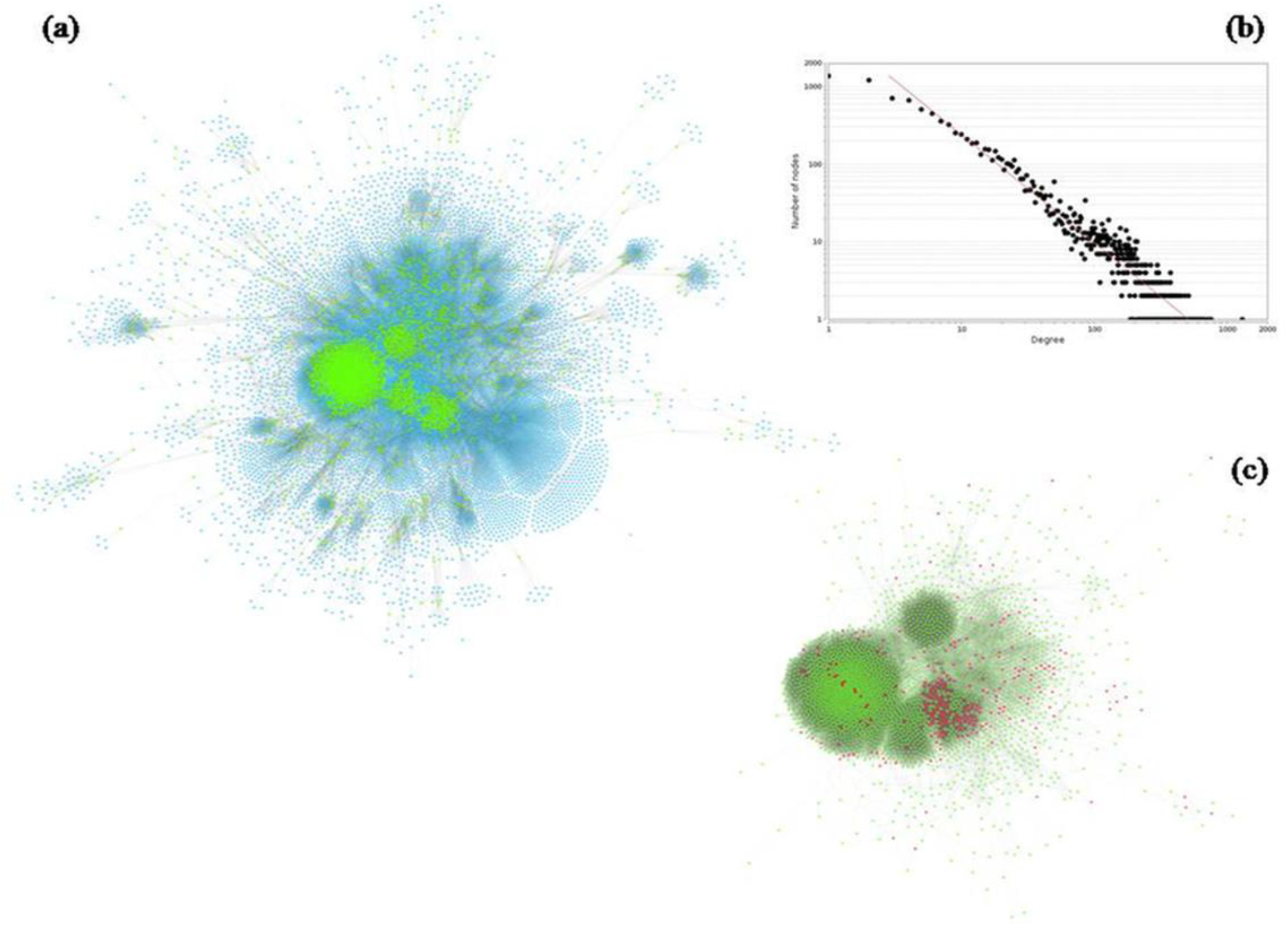
(a) Tea leaf protein-protein interaction network (TeaLIPIN). Green coloured nodes represent the identified key proteins in overall network. (b) Power-law fit on the degree distribution of TeaLIPIN. (c) A subnetwork of TeaLIPIN consisting of key proteins (green colour) and their interactions among themselves. Transcription factors in the key proteins are highlighted with red colour.

### Exploring the modular architecture of TeaLIPIN

Various complex biological processes are carried-out within the cell by multiple interactions among selected group of proteins, identification of functional modules in PPI network is an important aspect for knowledge discovery in biological systems. MCODE assisted clustering analysis of entire TeaLIPIN resulted in 207 modules consisting of total 4031 transcripts [S4 Supplementary File]. Based on total number of nodes in a cluster, we selected top 10 modules for pathways enrichment and found that 644 transcripts are encoding to 284 unique KO ids. Pathway enrichment analysis of these top-10 modules resulted in the identification of four enriched pathways related to genetic information processing that are ribosome, proteasome, spliceosome and ubiquitin mediated proteolysis and one enriched pathway related to plant pathogen interaction under organismal system category.

Ribosome pathway is found to be highly enriched (86 proteins from module_1, 104 proteins from module_2, 108 proteins from module_4 and 64 proteins from module_10) followed by Spliceosome pathway (27 proteins in module_5, 29 proteins in module_8 and 22 proteins in module_13), Proteasome pathway (43 proteins in module_3) and Ubiquitin mediated proteolysis (16 proteins in module_20). Plant pathogen interaction pathway is found to be enriched in module_30 with 6 proteins in environmental adaption under organismal systems category. Plant pathogen interaction pathway is important for recognition of pathogen attacks by pattern recognition receptors or effectors triggered immunity (Bellincampi et al. 2014). While exploring the details of proteins in this category, it is observed that TeaL_175716 and TeaL_159132 act as heat shock proteins that help in enhancing the synthesis of required proteins in stress condition (Kim and Yenari 2017), TeaL_176108 and TeaL_161500 are known to have role in MAP kinase signalling to encounter the abiotic stress (Zhu 2016) and the other two proteins TeaL_153931 and TeaL_189530 are leucine rich repeats that along with different receptor kinases possibly play regulatory roles in signal transduction and propagation during biotic stresses (Jayaswall et al. 2016).

### Identification of key proteins in TeaLIPIN and their TF characterization

During network analysis it was found that some nodes are in solitary clusters, thus largest component was selected for further analysis that consisted of 10,922 nodes and 1,97,309 interactions. To identify the key proteins in the TeaLIPIN, this largest component was subjected for 10,000 Erdős-Rényi (G_n,m_ type) model based realizations of random networks. Thereafter, by calculating the z-scores of all the protein nodes on the basis of three crucial centrality measures (degree, betweenness and eigenvector), a total of 2,931 proteins were found to have *p*-value less than 0.01 and are termed as key proteins in the rest of this manuscript (Fig. 5). Further analysis of key proteins revealed that 1,523 proteins have important role in the regulation of 276 pathways and 270 proteins are in 34 transcription factor families, among these 150 are uniquely associated with 30 TF families [Supplementary Table S2]. Out of 270 transcription factors, highest numbers are present in S1Fa-like TF (112 proteins) family followed by WRKY (89 proteins) and NAC (79 proteins) TF families (Fig. 5 (b)). In the following, we discuss about our findings in some of the TF families.

S1Fa-like transcription factors are activated by proteolytic processing in plants to regulate gene expression in response to various stimuli (Yao et al. 2017). It is found that among 112 key proteins that are related to S1Fa-like TF family, 8 proteins are unique to S1Fa-like TF family. Of these 8 proteins, TeaL_113200 has the highest degree (105) followed by TeaL_219351 (62 degree). TeaL_113200 is a MDIS1-interacting receptor like kinase and has important role in breaking reproductive isolation barrier (Wang et al. 2016). TeaL_219351 is CDK7 (cyclin-dependent kinase 7) having KO id K02202 under genetic information processing and is known to trigger the translational regulations by intricate feedback regulations of particular stress conditions (Merchante et al. 2015). Out of 89 proteins in WRKY TF family, 15 proteins (TeaL_208517; 109 interactions, TeaL_29412; 65 interactions, TeaL_10183; 61 interactions, TeaL_251204; 60 interactions etc.) are specific to this category. WRKY TFs have crucial role in molecular regulation during stress caused by pathogen attack (Mao et al. 2011), and are known to play important regulatory roles in various stress tolerance processes in plants by regulating the expression of other defense related genes that is also reported in tea (Wu et al. 2016). In NAC TF family, out of 79 proteins 6 are unique to this category that include TeaL_160350 with highest degree 55 followed TeaL_8246 with degree 34. NAC TFs are known to manage ROS load during both biotic and abiotic stress in rice (Fang et al. 2015) and may also be helpful in tea stress conditions.

Among other TF categories, we have found 21 bHLH (TeaL_191930; 422 interactions, TeaL_255207; 324 interactions, TeaL_188079; 275 interactions etc.) and 9 MYB/MYB_related (TeaL_3029; 168 interactions, TeaL_8162; 66 interactions etc.) transcription factors. Different complexes of MYB/bHLH transcription factors regulate several distinct processes in a cell which include hormonal signaling, circadian clock, cell wall synthesis, cell death, biotic and abiotic stress responses, and biosynthesis of metabolites for specific processes. In several other cases, physical interactions of bHLHs take place with MYBs in order to regulate the function based on cell demand at particular condition (Stracke et al. 2001; Seo and Mas 2014; Pireyre and Burow 2015). Also, Whirly transcription factors are known to have crucial roles in regulating the expression of genes during stress conditions (biotic and abiotic) by reducing ROS accumulation (Zhao et al. 2018). In this work, we have found 17 Whirly transcription factors (TeaL_121652; 297 interactions, TeaL_209827; 163 interactions, TeaL_1629; 145 interactions, TeaL_15392; 123 interactions) that might have regulatory role for stress tolerance in tea also.

### *In-silico* validation of transcripts involved in TeaLIPIN

In order to validate the existence of network proteins in the genome, all the 11,206 transcripts were aligned with draft tea genome. A total of 8,859 transcripts successfully mapped to 3,578 unique fragments of draft genome. Additionally, among all identified key proteins (270 in number) encoding transcription factors, 136 proteins were successfully mapped to available protein data of *Camellia sinensis* at NCBI and 234 transcripts were mapped to genome with 192 unique fragments [S5 Supplementary File]. Hence, these proteins might be thought of as the novel key candidates and may be studied in detail for understanding the regulatory processes related to biotic and abiotic stresses.

## Summary

To the best of our knowledge this is the first work in tea where an interolog based PPI network is reported. The methodology developed to construct this network utilizes the publicly available transcriptomes without relying on reference assembly or the annotated genome. Investigations based on protein-protein network interactions by means of wet-lab experiments require immense efforts, financial aid and time consumption; therefore identification of high-confidence PPI interactions through computational methods is gaining much recognition in recent times. Although complex transcriptional mechanism can be illustrated using comparative analysis by *in-silico* tools but the identification of key proteins in a particular pathway and determining their interactors with certain confidence require in-depth computational analysis that can enhance the success propensity as well as reduce the financial burden (of hit-and-trial) *in-vivo* or *in-vitro* experiments.

Tea is one of the most popular and most ancient therapeutic beverages and is obtained from newly born young leaves that continuously experience attack of various biotic and abiotic stresses. To capture the holistic response mechanisms by the tea leaf, our input dataset was prepared by developing a global assembly of tea leaf transcripts from six publically available RNA-Seq datasets comprising three biotic stress conditions and three abiotic stress conditions. This assembly consisting of 2,61,695 final assembled transcripts was firstly annotated using various bioinformatics protocols for functional characterization, specifically pathway assignment and transcription factor identification. To gain insights of response mechanisms that are generally activated via a web of interactions among proteins instead of a single one, we then used the annotated transcripts to develop a draft protein-protein interactions network in a tea leaf (TeaLIPIN) consisting of 11,208 nodes and 1,97,820 edges. To enhance the tenacity of interactions among proteins, we relied only on the experimentally known protein interactions in 14 plant species. This is probably the largest number of species used for inferring an interologous PPI network of any plant so far.

By comparing the three important network metrics of TeaLIPIN with its 10,000 random realizations (G_n,m_ type), we have identified 2,931 key candidates related to growth, development and stress tolerance. A total of 1,523 proteins are successfully assigned to various important regulatory pathways and 270 proteins are predicted to be potential transcription factors using in-house developed hidden Markov models, out of these 150 are found to be uniquely present in any one of the 30 transcription factor families and 60 are present as TFs of tea in the NCBI’s Protein database. Additionally, MCODE clustering of the final network resulted in 207 clustering modules. Based on cluster size, top 10 modules were studied for their functional enrichment. This division of TeaLIPIN into biological meaningful modules may aid to the automated detection of protein interactors and prediction of biological processes to help us uncover the complex circuitry design in the cell of tea leaf.

Moreover, in order to further enhance the confidence in the annotated transcripts from developed assembly these were mapped with the draft genome of tea (Xia et al. 2017) that has been recently reported and could successfully map 243,885 out of 261,695 final assembled transcripts. Interestingly, all the key proteins reported here are successfully mapped to the tea draft genome.

In the future, TeaLIPIN proposed in this work may be used to identify the potential therapeutic targets and the key candidates may be utilized for crop improvement by plant breeding methods. We believe that the computational framework developed in this study can be easily adopted to build useful knowledgebase of networks at high scale of complexity for other plant species too.

## Acknowledgments

We thank Central University of Himachal Pradesh for providing the computational infrastructure.

## Authors’ Contributions

VS* conceptualized the research framework and supervised the work. GS and VS^†^ performed the computational studies. All the authors analyzed the data and interpreted results. GS and VS* wrote and finalized the manuscript.

## Supplementary Materials

**S1 Supplementary File:** Sequences of all the final assembled transcripts

**S2 Supplementary File:** Annotation details of assembled sequences.

**S3Supplementary File:** TeaLIPIN interactions

**S4 Supplementary File:** Detailed list of all the identified functional modules, pathways analysis of key proteins and transcription factors identified in key proteins.

**S5 Supplementary File:** Mapping of TeaLIPIN proteins on draft genome of tea and available proteomic data at NCBI.

## References

1. Altschul SF, Gish W, Miller W, et al (1990) Basic local alignment search tool. J Mol Biol 215:403–410

2. Apweiler R, Bairoch A, Wu CH, et al (2004) UniProt: the universal protein knowledgebase. Nucleic Acids Res 32:D115–D119

3. Bader GD, Hogue CW V (2003) An automated method for finding molecular complexes in large protein interaction networks. BMC Bioinformatics 4:2

4. Bellincampi D, Cervone F, Lionetti V (2014) Plant cell wall dynamics and wall-related susceptibility in plant–pathogen interactions. Front Plant Sci 5:228

5. Berardini TZ, Reiser L, Li D, et al (2015) The Arabidopsis information resource: making and mining the “gold standard” annotated reference plant genome. genesis 53:474–485

6. Du Z, Zhou X, Ling Y, et al (2010) agriGO: a GO analysis toolkit for the agricultural community. Nucleic Acids Res 38:W64–W70

7. Erdos P, Rényi A (1960) On the evolution of random graphs. Publ Math Inst Hung Acad Sci 5:17–60

8. Fang Y, Liao K, Du H, et al (2015) A stress-responsive NAC transcription factor SNAC3 confers heat and drought tolerance through modulation of reactive oxygen species in rice. J Exp Bot 66:6803–6817

9. Geisler-Lee J, O’Toole N, Ammar R, et al (2007) A predicted interactome for Arabidopsis. Plant Physiol 145:317–329

10. Grabherr MG, Haas BJ, Yassour M, et al (2011) Full-length transcriptome assembly from RNA-Seq data without a reference genome. Nat Biotechnol 29:644

11. Huang DW, Sherman BT, Lempicki RA (2008) Bioinformatics enrichment tools: paths toward the comprehensive functional analysis of large gene lists. Nucleic Acids Res 37:1–13

12. Jayaswall K, Mahajan P, Singh G, et al (2016) Transcriptome analysis reveals candidate genes involved in blister blight defense in tea (*Camellia sinensis* (L) Kuntze). Sci Rep 6:30412

13. Jeong H, Mason SP, Barabási A-L, Oltvai ZN (2001) Lethality and centrality in protein networks. Nature 411:41

14. Jin J, Tian F, Yang D-C, et al (2016) PlantTFDB 4.0: toward a central hub for transcription factors and regulatory interactions in plants. Nucleic Acids Res gkw982

15. Joy MP, Brock A, Ingber DE, Huang S (2005) High-betweenness proteins in the yeast protein interaction network. Biomed Res Int 2005:96–103

16. Kanehisa M, Furumichi M, Tanabe M, et al (2016) KEGG: new perspectives on genomes, pathways, diseases and drugs. Nucleic Acids Res 45:D353–D361

17. Kim JY, Yenari M (2017) Heat Shock Proteins and the Stress Response. In: Primer on Cerebrovascular Diseases (Second Edition). Elsevier, pp 273–275

18. Li C-F, Xu Y-X, Ma J-Q, et al (2016) Biochemical and transcriptomic analyses reveal different metabolite biosynthesis profiles among three color and developmental stages in ‘Anji Baicha’(Camellia *sinensis*). BMC Plant Biol 16:195

19. Li W, Godzik A (2006) Cd-hit: a fast program for clustering and comparing large sets of protein or nucleotide sequences. Bioinformatics 22:1658–1659

20. Li X, Lin Y, Zhao S, et al (2018) Transcriptome changes and its effect on physiological and metabolic processes in tea plant during mechanical damage. For Pathol e12432

21. Mao G, Meng X, Liu Y, et al (2011) Phosphorylation of a WRKY transcription factor by two pathogen-responsive MAPKs drives phytoalexin biosynthesis in Arabidopsis. Plant Cell tpc-111

22. Merchante C, Brumos J, Yun J, et al (2015) Gene-specific translation regulation mediated by the hormone-signaling molecule EIN2. Cell 163:684–697

23. Namita P, Mukesh R, Vijay KJ (2012) *Camellia Sinensis* (green tea): A review. Glob J Pharmacol 6:52–59

24. Newman, M. E. (2008). The mathematics of networks. The new palgraveencyclopedia of economics, 2:1–12.

25. Patel RK, Jain M (2012) NGS QC Toolkit: a toolkit for quality control of next generation sequencing data. PLoS One 7:e30619

26. Paul A, Jha A, Bhardwaj S, et al (2014) RNA-seq-mediated transcriptome analysis of actively growing and winter dormant shoots identifies non-deciduous habit of evergreen tree tea during winters. Sci Rep 4:5932

27. Pireyre M, Burow M (2015) Regulation of MYB and bHLH transcription factors: a glance at the protein level. Mol Plant 8:378–388

28. Seo PJ, Mas P (2014) Multiple layers of posttranslational regulation refine circadian clock activity in Arabidopsis. Plant Cell tpc-113

29. Shi J, Ma C, Qi D, et al (2015) Transcriptional responses and flavor volatiles biosynthesis in methyl jasmonate-treated tea leaves. BMC Plant Biol 15:233

30. Sievers F, Wilm A, Dineen D, et al (2011) Fast, scalable generation of high-quality protein multiple sequence alignments using Clustal Omega. Mol SystBiol 7:539

31. Smoot ME, Ono K, Ruscheinski J, et al (2010) Cytoscape 2.8: new features for data integration and network visualization. Bioinformatics 27:431–432

32. Stracke R, Werber M, Weisshaar B (2001) The R2R3-MYB gene family in *Arabidopsis thaliana*. CurrOpin Plant Biol 4:447–456

33. Szklarczyk D, Franceschini A, Wyder S, et al (2014) STRING v10: protein–protein interaction networks, integrated over the tree of life. Nucleic Acids Res 43:D447–D452

34. Thanasomboon R, Kalapanulak S, Netrphan S, Saithong T (2017) Prediction of cassava protein interactome based on interolog method. Sci Rep 7:17206

35. Wang T, Liang L, Xue Y, et al (2016) A receptor heteromer mediates the male perception of female attractants in plants. Nature 531:241

36. Wu Z-J, Li X-H, Liu Z-W, et al (2016) Transcriptome-wide identification of *Camellia sinensis* WRKY transcription factors in response to temperature stress. Mol Genet genomics 291:255–269

37. Xia E-H, Zhang H-B, Sheng J, et al (2017) The tea tree genome provides insights into tea flavor and independent evolution of caffeine biosynthesis. Mol Plant 10:866–877

38. Yao S, Deng L, Zeng K (2017) Genome-wide in silico identification of membrane-bound transcription factors in plant species. PeerJ 5:e4051

39. Ye J, Fang L, Zheng H, et al (2006) WEGO: a web tool for plotting GO annotations. Nucleic Acids Res 34:W293–W297

40. Yu H, Luscombe NM, Lu HX, et al (2004) Annotation transfer between genomes: protein–protein interologs and protein–DNA regulogs. Genome Res 14:1107–1118

41. Zhang Q, Cai M, Yu X, et al (2017) Transcriptome dynamics of *Camellia sinensis* in response to continuous salinity and drought stress. Tree Genet Genomes 13:78

42. Zhang S, Jin G, Zhang X, Chen L (2007) Discovering functions and revealing mechanisms at molecular level from biological networks. Proteomics 7:2856–2869

43. Zhao S-Y, Wang G-D, Zhao W-Y, et al (2018) Overexpression of tomato WHIRLY protein enhances tolerance to drought stress and resistance to *Pseudomonas solanacearum* in transgenic tobacco. Biol Plant 62:55–68

44. Zhou Y, Liu Y, Wang S, et al (2017) Molecular cloning and characterization of galactinol synthases in Camellia sinensis with different responses to biotic and abiotic stressors. J Agric Food Chem 65:2751–2759

45. Zhu G, Wu A, Xu X-J, et al (2016) PPIM: a protein-protein interaction database for maize. Plant Physiol 170:618–626

46. Zhu J-K (2016) Abiotic stress signaling and responses in plants. Cell 167:313–324

47. Zhu P, Gu H, Jiao Y, et al (2011) Computational identification of protein-protein interactions in rice based on the predicted rice interactome network. Genomics Proteomics Bioinformatics 9:128–137

